# Magnetic Activation of Spherical Nucleic Acids for the Remote Control of Synthetic Cells

**DOI:** 10.1101/2024.08.21.608917

**Authors:** Ellen Parkes, Assala Al Samad, Giacomo Mazzotti, Charlie Newell, Brian Ng, Amy Radford, Michael J. Booth

## Abstract

The advancement of synthetic cells as drug delivery devices hinges on the development of targeting strategies, in particular the controlled synthesis of biomolecules in-situ using a deeply penetrative stimulus. To address this, we have designed spherical nucleic acids comprising DNA promoter sequences decorating magnetic nanoparticle cores. By harnessing the heat dissipated from magnetic hyperthermia (a clinically-approved anticancer therapy) we tightly controlled cell-free protein synthesis. We then deployed a tissue phantom that is impenetrable by current activation methods to demonstrate the potential of this technology for the remote control of synthetic cells using deeply tissue-penetrating magnetic fields. This paves the way for targeting and controlling the in-situ synthesis of biomolecules deep within the body.

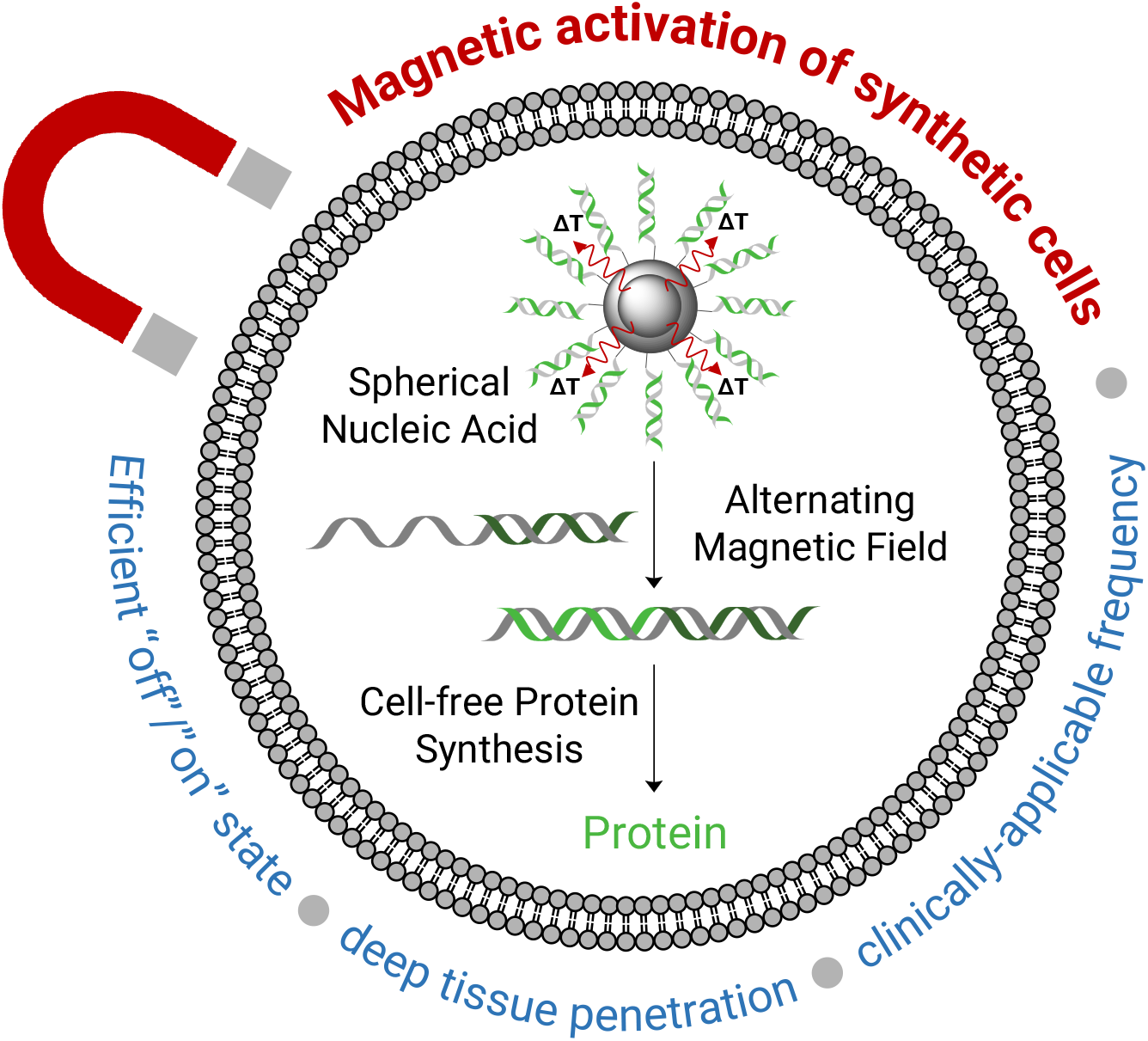

## Introduction

In recent years the complexity of reprogramming living cells has driven research efforts into constructing synthetic mimics using modular parts. Biomimicry of enzymatic reactions and stimuli-responses have been achieved in synthetic cells; non-living compartments capable of imitating the behaviour, function, and structure of living cells (*1, 2*). The majority of synthetic cells comprise a cell-free protein synthesis (CFPS) system bounded by a lipid membrane, programmed according to DNA templates that are specific to the desired biological activity (*3*). Notably, synthetic cells can accommodate non-biological components to attain new functionalities that are not accessible to living cells, which significantly increases their technological value by opening up applications such as novel drug delivery systems and bioreactors (*2, 4*).

A prerequisite for the application of synthetic cells to biology and medicine is the ability to control their activity, reducing off-target effects and improving therapeutic potential (*5*). Modulating CFPS at the DNA or mRNA level has been achieved using small-molecule-sensitive transcription factors and riboswitches, though approaches of these kind are limited in synthetic cells due to the difficulty of small molecules traversing the membrane (*6*–*8*). By using an external stimulus, remote regulation of internal CFPS can be achieved in synthetic cells with a level of spatial precision not accessible to small-molecule techniques (*5, 9*). The majority of research in this area currently focuses on using ultraviolet (UV) light, which poses cytotoxicity issues and does not penetrate tissue, hindering its use in therapeutic applications (*5*). Beyond this, temperature has been explored as an external stimulus to control CFPS and small molecule release with RNA thermometers, albeit at temperatures outside of the biologically useful range (*10*). The ideal external stimulus for controlling CFPS within synthetic cells is alternating magnetic fields (AMFs), which are deeply tissue-penetrating (>10 cm) and biologically benign (*11, 12*). Magnetic nanoparticles dissipate thermal energy to the surrounding medium as a result of Brownian and Néel fluctuations in the presence of an AMF (*13*). Brownian relaxation is described as the physical rotation of the particle, whereas Néel relaxation is described as the rotation of the magnetic moment within the particle. If the applied AMF is strong enough to rotate the particle itself or to displace the magnetic moment from the preferred orientation, then relaxation back to equilibrium releases thermal energy to the surroundings (*14*). This localised heating, termed magnetic hyperthermia, is clinically-approved to ablate malignant tumours by exploiting the differing heat tolerance between tumour and healthy cells, whereby healthy cells can survive above physiological temperature for longer (*15*). Magnetic hyperthermia has also been investigated for rupturing lipid bilayers, but it demands high frequencies that are neither clinically tolerable nor compatible with current medical AMF generators (*16–19*). Additionally, the disruption of bilayers limits their potential use with synthetic cells and CFPS.

We have designed a magnetically controllable CFPS system for use in synthetic cells by taking inspiration from spherical nucleic acids (SNAs), a delivery platform comprising nucleic acids that are radially arranged around a nanoparticle core (*20*). We have synthesised SNAs with magnetic cores to harness the heat dissipated from magnetic hyperthermia and release a T7 promoter sequence, enabling the recovery of an otherwise inactive DNA template and the subsequent spatiotemporal control of its activity. This mechanism has previously been applied to measure the surface temperature of nanoparticles and to release DNA *in-vivo*, but required high electromagnetic frequencies (*11, 21, 22*). In addition, these previous studies did not control the activity and downstream function of the attached DNA. Through nanoparticle engineering and the development of a novel SNA purification method, we were able to activate gene-expressing synthetic cells within tissue phantoms using clinically tolerable and FDA/EMA approved frequencies (100 kHz). This technology paves the way for the application of synthetic cells as a widespread therapeutic modality and realises their potential as on-demand drug delivery devices with enhanced tissue penetration.

## Results

### Magnetic nanoparticle engineering

To create our magnetically-activated spherical nucleic acids, we designed the central scaffold to consist of silica-encapsulated iron oxide nanoparticles (IONPs@SiO_2_) (Fig. 1). We synthesised IONPs by thermally decomposing iron(III) acetylacetonate in the presence of oleylamine and benzyl ether (*23, 24*). The hydrophobic oleylamine-capped IONPs were of uniform shape and narrow size distribution, as confirmed by transmission electron microscopy (TEM) with a diameter of *D*_TEM_ = 6.4 ± 1.0 nm (Fig. 1A). Further structural information was obtained by X-ray diffraction (XRD), whereby the oleylamine-capped IONPs were crystalline and phase pure with the position and relative intensity of the diffraction peaks matching that of magnetite (RUFF ID: R061111) (Fig. S1A) (*25*). The crystallite size calculated from the Scherrer equation at *D*_XRD_ = 6.2 nm was comparable to that determined by statistical analysis of the TEM images, indicating that each particle was a single crystal (*26*). Dynamic Light Scattering (DLS) was utilised to investigate the hydrodynamic size of the nanoparticles and proved larger than the crystallite size at *D*_DLS_ = 8.4 nm, taking into account the fluid interactions and adsorbed surfactant (Table 1). The hysteresis curve measured at 300 K determined that the oleylamine-capped IONPs were superparamagnetic at room temperature with zero coercivity, in-turn confirming that the net magnetisation in the absence of an external field was zero (Fig. S1B). Their saturation magnetisation plateaued at 56.0 emu g_Fe_^-1^ whilst their magnetic susceptibility (χmax) equalled 0.049 emu g^-1^ Oe^-1^ at 68.5 Oe, the former property influences the thermal energy dissipated in an AMF with high saturation magnetism values resulting in greater inductive heating (*27*).

**Table 1.** Nanoparticle characterisation. Summary of the hydrodynamic diameter (analysis in number), polydispersity index (PDI) and ζ-potential, as determined by DLS with the mean calculated from n = 3 repeat measurements.

| Nanoparticle | Hydrodynamic diameter (nm) | PDI | ζ-potential (mV) |
| --- | --- | --- | --- |
| Oleylamine-capped IONPs | 8.4 | 0.43 | N/A |
| NH<sub>2</sub>-modified IONPs@SiO<sub>2</sub> | 36.7 | 0.38 | 17.8 |
| DBCO-modified IONPs@SiO<sub>2</sub> | 142.0 | 0.20 | -13.9 |
| dsDNA T7-modified IONPs@SiO<sub>2</sub> (SNA) | 106.3 | 0.33 | -36.6 |

To obtain biocompatible particles with a readily modifiable surface chemistry, we coated the IONPs in silica (SiO_2_) (Fig. 1B). The reverse microemulsion route was chosen for enhanced morphological control and reduced yields of by-product SiO_2_ with no IONP core (*28*). Briefly, the oleylamine-capped IONPs underwent a ligand exchange with the surfactant IGEPAL-*co*-520 in cyclohexane, prior to the addition of concentrated aqueous ammonium hydroxide that nucleated the tetraethylorthosilicate precursor within the as-formed aqueous domains, forming SiO_2_ shells around single IONP cores. In the final step (3-aminopropyl)triethoxysilane) was injected to modify the surface of the IONPs@SiO_2_ with reactive amine (NH_2_) groups. The NH_2_-modified IONPs@SiO_2_ were characterised by TEM, revealing monodisperse particles with a diameter of *D*_TEM_ = 20.9 ± 1.8 nm and no observed by-product SiO_2_ (Fig. 1B). DLS was utilised to investigate the hydrodynamic size after the SiO_2_ encapsulation, which increased to *D*_DLS_ = 36.7 nm. The zeta (ζ) potential further indicated the addition of protonatable NH_2_ groups with a measured charge of ζ = +17.8 mV (Table 1). A copper-free click handle was installed on the surface of the NH_2_-modified IONPs@SiO_2_ by coupling to an *N*-hydroxysuccinimide (NHS) ester-modified dibenzocyclooctyne (DBCO). The success of the reaction was tracked by ultraviolet-visible (UV-vis) spectroscopy and the absorbance of DBCO at its excitation maximum wavelength of 309 nm (Fig. 1C). The Beer Lambert Law was used to decipher the concentration of DBCO at a molar extinction coefficient of ε = 12,000 M^-1^cm^-1^ (309 nm) (Eq. S5,6) (*29*). The particle concentration was determined by approximating individual nanoparticles as spherical in shape and relating the volume and area extrapolated from TEM analysis to known densities of magnetite and silica (Eq. S1–4) (*30–32*). The resulting number of DBCO molecules decorating each core-shell nanoparticle equaled 3669. DLS measurements inferred the successful modification by DBCO with an increase in hydrodynamic diameter from *D*_DLS_ = 36.7 nm for NH_2_-modified IONPs@SiO_2_ compared with *D*_DLS_ = 142.0 nm for DBCO-modified IONPs@SiO_2_ and further a shift in surface charge from ζ = +17.8 mV to ζ = -13.9 mV (Table 1), consistent with the change in surface chemistry.

**Fig. 1.**
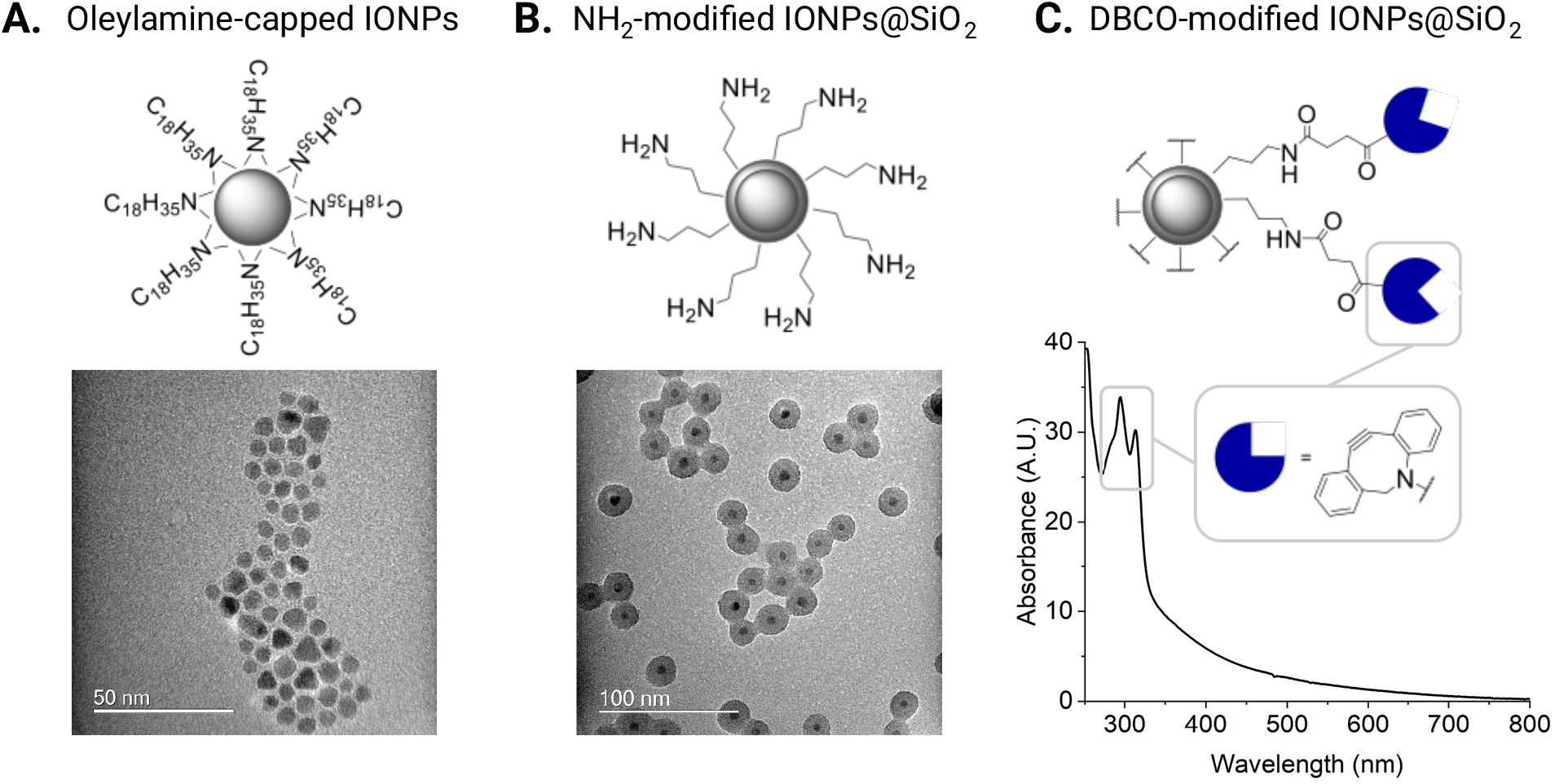
Silica-encapsulated iron oxide nanoparticle construction and their modification with the DBCO click handle. **(A)** Schematic and TEM image of oleylamine-capped IONPs synthesised by thermal decomposition. **(B)** Schematic and TEM image of NH_2_-modified IONPs@SiO_2_ synthesised by reverse-microemulsion. **(C)** Schematic and UV-Vis spectrum of DBCO-modified IONPs synthesised by NHS ester-coupling, highlighting the characteristic DBCO absorption.

### Spherical nucleic acid synthesis

In order to form the spherical nucleic acids (SNAs), we used copper-free strain-promoted azide– alkyne cycloaddition to conjugate azide-modified double-stranded (ds) DNA, comprising the T7 promoter strand, to the DBCO-modified IONPs@SiO_2_ (Fig. 2A,B). In this system, the “bottom” T7 promoter strand was covalently attached to the nanoparticles with the “top” T7 promoter strand hybridised to the bottom strand. The bottom strand of the T7 promoter was sourced with a 5’-azide group to enable conjugation to the surface of the nanoparticles. This azide-modified bottom T7 promoter strand was annealed to an unmodified top T7 promoter strand and analysed by poly(acrylamide) gel electrophoresis (PAGE) (Fig. S2). The SNA synthesis was carried out by freezing the azide-modified dsDNA in the presence of DBCO-modified IONPs@SiO_2_ and low concentrations of salt (*33*). The freeze-directed construction of SNAs takes advantage of ice crystals gradually forming, concentrating the reagents into “micro-pockets” and alleviating steric and electrostatic hindrances, thereby encouraging SNA formation (*33, 34*). We identified high concentrations of DNA electrostatically-bound to the nanoparticles (alongside the formation of covalently-bound DNA) with conventional SNA synthesis methods. This was detrimental to controlling the activity of the attached DNA, as the electrostatically-bound DNA leached off in biological conditions (Fig S3A). To mediate this, we developed a purification method to produce SNAs decorated solely with covalently-bound DNA. Our purification involved pulling electrostatically-bound DNA from the surface of the nanoparticle using an electric current that was applied through an agarose gel; the covalently-constructed SNA did not travel through the gel and was then extracted through brief sonication of the excised well in water (Fig. 2C). We found that this purification step was essential to rendering the SNA inactive in the absence of the AMF (Fig. S3A).

**Fig. 2.**
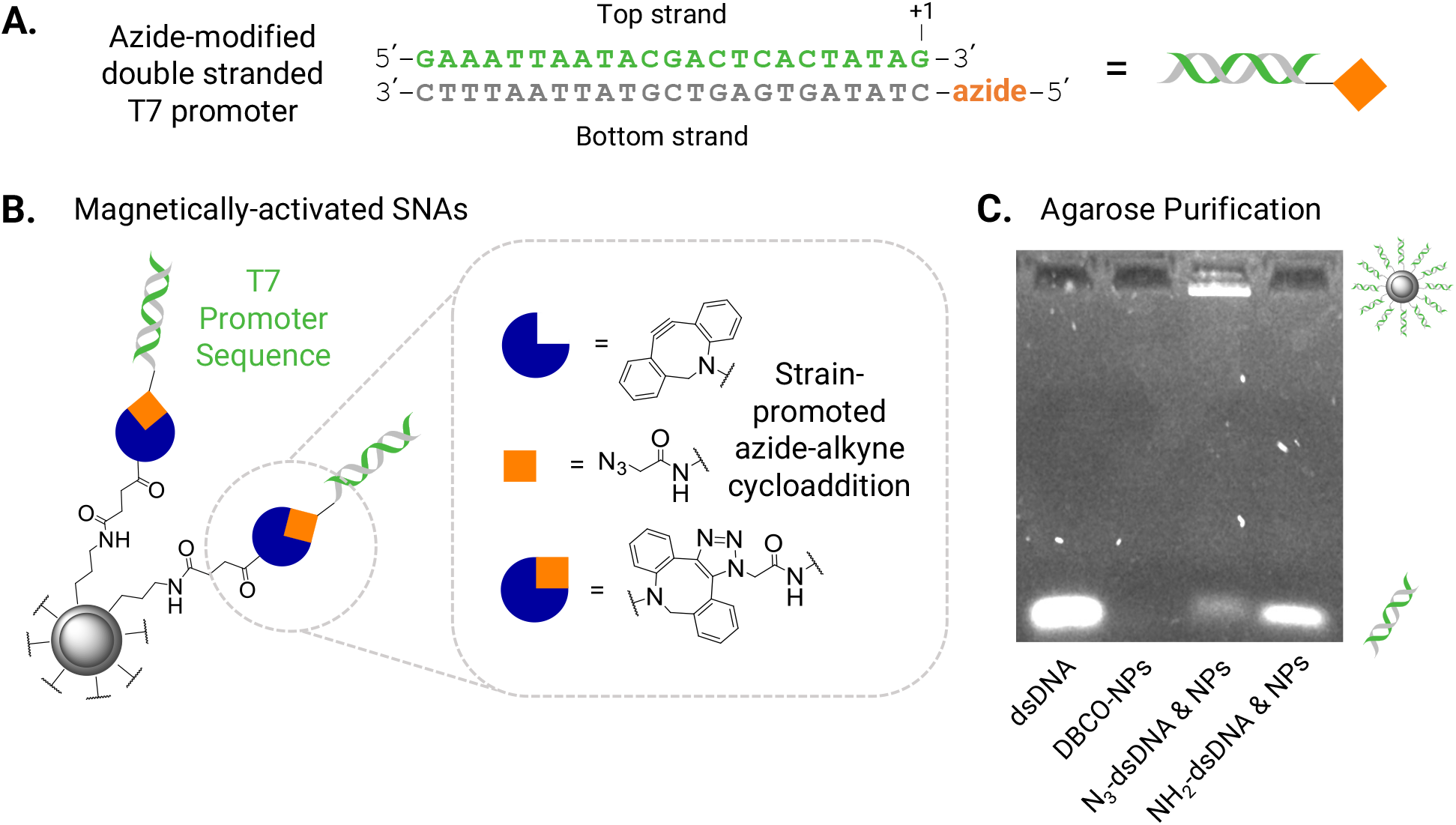
Spherical nucleic acid synthesis and purification. **(A)** Schematic showing the top and bottom (azide-modified) T7 promoter strands. The transcription initiation site (+1) is marked. **(B)** Schematic detailing the covalent attachment of the T7 promoter strands to the surface of the nanoparticle to form the magnetically-activated SNAs. The azide-modified, double stranded T7 promoter enabled copper-free click chemistry to the DBCO-modified nanoparticles. **(C)** Agarose gel showing the SNA purification method to remove electrostatically-bound DNA from the surface of the nanoparticles using an electric current. The covalently-constructed SNA (with the azide-modified DNA) did not travel through the gel and was then extracted through brief sonication of the excised well in water. DNA that was not covalently attached to the surface of the nanoparticles was pulled-off by applying an electric current to the agarose gel and can be seen to travel through the gel at the same speed as the free DNA and amine-only modified dsDNA incubated with the nanoparticles.

The resulting magnetically-activated SNAs remained colloidally stable as determined by the highly negative ζ-potential induced by the phosphate DNA backbone, decreasing to ζ = -36.6 mV (Table 1). The loading of the T7 promoter was determined by denaturing the dsDNA attached to the IONPs@SiO_2_ at 95°C for 5 min in urea-containing dye and comparing the amount of released DNA to a calibration curve of known concentration on an agarose gel (Fig. S4 and detailed in the SI). The calculated concentration of attached T7 promoter strands was related to the previously calculated nanoparticle concentration to give an estimated loading of 18 dsDNA strands per nanoparticle, which is in line with literature values for SNAs comprising dsDNA (*33*).

### Magnetic activation of cell-free protein synthesis

To control CFPS with magnetism, we synthesised an inactive DNA template encoding the fluorescent protein mNeonGreen (mNG) with a single-stranded 3’ overhang of the bottom T7 promoter sequence (on the template strand) and the top T7 promoter sequence (on the non-template strand) missing, taking advantage of T7 RNA polymerase only transcribing from a dsT7 promoter sequence (*35*). To achieve this, we synthesised two mNG DNA templates by polymerase chain reaction (PCR), one with and one without the dsDNA T7 promoter region, and then used lambda exonuclease to generate the respective single-stranded versions to anneal together (Fig. S5A,B). A 5’-phosphorylated primer was used for the mNG non-template strand with the promoter region and the mNG template strand without the promoter region. This enabled the digestion of the individual phosphorylated DNA strands by lambda exonuclease. The intact mNG template single strand with the promoter region and the intact mNG non-template single strand without the promoter region were annealed to produce the inactive mNG DNA template (Fig. S5A,B). To calculate the loss of activity of the inactive template, we used a commercial CFPS system (PURExpress) and measured the fluorescence intensity of expressed mNG protein after incubation for 3 h. Compared with the full template, containing the intact dsDNA T7 promoter region, the inactive template had 6% activity (Fig. 3B). Whereas using no DNA template gave 2% activity, compared with the full template. We explored further the influence of the IONPs@SiO_2_ alone on mNG expression and saw minimal difference in the activity of PURExpress and the fluorescence intensity of mNG expression with and without the nanoparticles present (Fig. S3B).

**Fig. 3.**
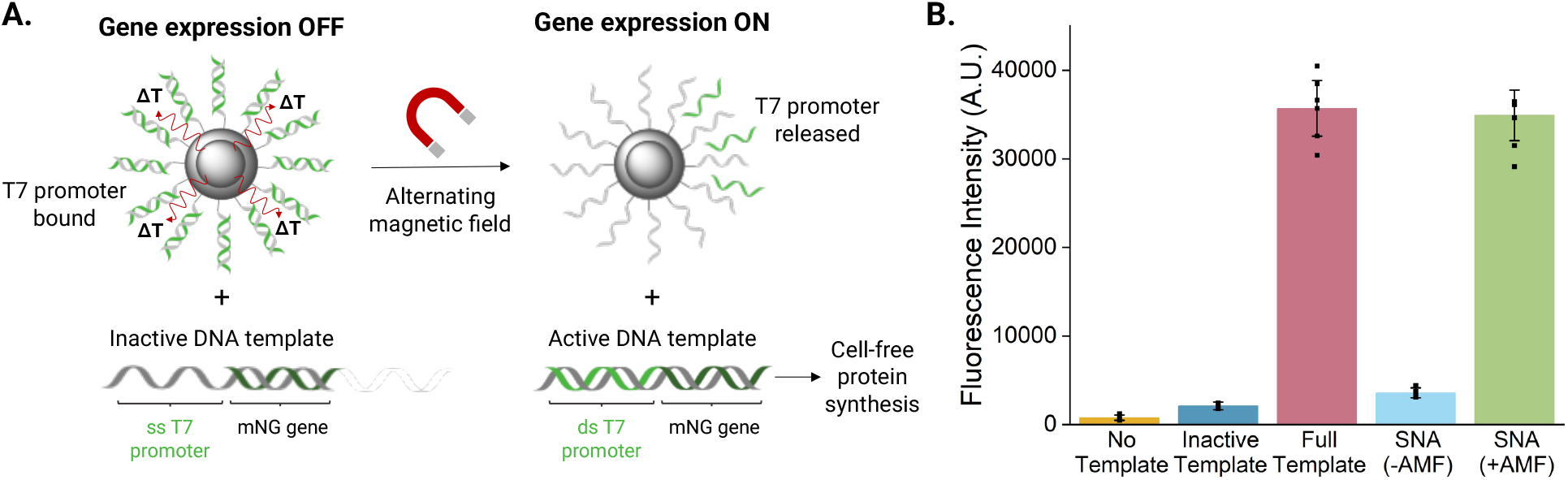
Controlling cell-free protein synthesis with an alternating magnetic field. **(A)** Schematic detailing the release of the T7 promoter ‘ top’ sequence from the SNA surface as a result of inductive heating and exposure to an AMF, and the subsequent hybridisation of the promoter sequence to the single-stranded T7 promoter of the inactive template, restoring the dsDNA T7 promoter and activating expression of the encoded fluorescent mNG protein. Note: promoter (23 bp) and gene (∼700 bp) length are not to scale. **(B)** Cell-free protein synthesis of mNG in the presence of magnetically-activated SNAs, with and without exposure to an AMF. Reactions that were not exposed to an AMF expressed minimal mNG, and their fluorescence intensity was comparable to the inactive DNA template only. Reactions exposed to an AMF expressed mNG and their fluorescence was consistent with reactions containing the full mNG template (dsDNA T7 promoter region present).

The inactive DNA template was combined with our magnetically-activated SNAs (containing the T7 promoter sequence) to control CFPS with an AMF (Fig. 3A). The localised thermal energy dissipated from the magnetic nanoparticles under an applied AMF was hypothesised to denature the conjugated dsDNA and release the hybridised T7 promoter (top strand). The newly released top strand would then be free to hybridise to the single-stranded T7 promoter of the inactive template, restoring the dsDNA T7 promoter and activating CFPS of the encoded fluorescent mNG protein. We applied an AMF through an 18-turn solenoid coil with parameters chosen to ensure operation within clinically tolerable frequencies (100 kHz) (*11, 19*). As such the AMF parameters selected were a field amplitude of 30 mT and a frequency of 103.4 kHz. The timeframe was mediated to ensure that the commercial CFPS system (PURExpress) did not experience global heating beyond physiological temperature (37°C) and was optimised to 25 min AMF exposure.

The magnetically-activated SNAs, inactive mNG template and CFPS system were exposed to an AMF, with a control experiment without an applied AMF. The resulting mNG expression was measured by fluorescence intensity after incubation for 3 h and compared with the inactive template alone (negative control) and the full mNG DNA template containing the dsDNA T7 promoter region (positive control), representing a theoretical 100% restoration of activity (Fig. 3B). In the control without an applied AMF, we observed only a minor increase in fluorescence compared with the inactive template (significant, p-value = 0.00025). In contrast, exposing the magnetically-activated SNAs to an AMF in the presence of the inactive mNG template and the CFPS system yielded a 98% recovery of expression compared with the full template (non-significant, p-value = 0.23), showing the tight regulation of biosynthesis *in-situ* and the selective “on” state only in the presence of the magnetic field. We carried out control reactions by replacing the magnetically-activated SNAs with the short dsDNA T7 promoter sequence along with the inactive template and exposed the reaction to an AMF. The lack of a significant increase in mNG expression determined that the selective “on” state was a result of inductive heating from magnetic hyperthermia and not simply the heat generated from the solenoid coil (Fig. S6).

### Magnetic activation of gene-expressing synthetic cells

To demonstrate magnetic-activation of CFPS-containing synthetic cells, we encapsulated the magnetically-activated SNAs, inactive mNG template, and CFPS system (PURExpress) inside giant unilamellar vesicles (GUVs). GUVs containing PURExpress were prepared as previously described, using egg phosphatidyl choline (egg-PC) and the inverted emulsion method (*3*). Briefly, an emulsion of the internal synthetic cell solution was prepared with a lipid-containing oil. The emulsion was then centrifuged through a phase transfer column, consisting of the outer buffer solution below the same lipid-containing oil. We encapsulated the PURExpress solution, magnetically-activated SNAs, the inactive DNA template encoding mNG, and Texas-Red-Dextran (TXR) to visualise the resulting GUVs. After preparation, the GUVs were exposed to an AMF (with a control not exposed to an AMF), incubated for 2.5 h and then imaged by fluorescence microscopy. The synthetic cells encapsulating the magnetically-activated SNAs and the inactive mNG template, but without exposure to an AMF showed minimal expression (median fluorescence intensity = 3.03 grey units), compared with the inactive template (median fluorescence intensity = 0.00 grey units), demonstrating an excellent “off” state (Fig. 4A,B). Excitingly, *in-situ* mNG expression inside the synthetic cells was fully recovered after exposure to an AMF, compared with the full template (median fluorescence intensity, 23.22 grey units vs 17.47 grey units). The enhanced mNG expression was observed only in the presence of the AMF, realising the remote control of synthetic cells with an AMF.

**Fig. 4.**
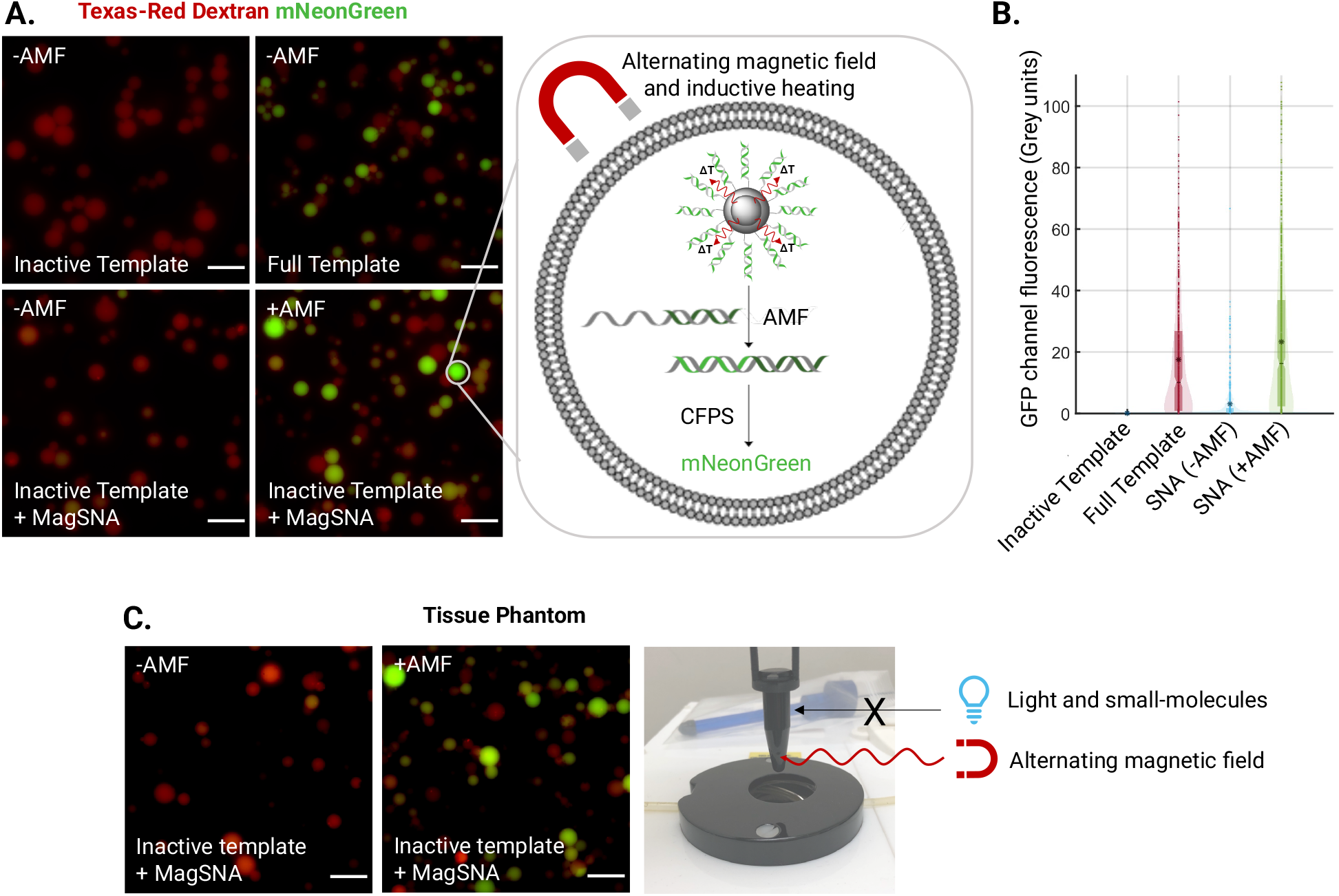
Controlling gene expression in synthetic cells with an alternating magnetic field. **(A)** Epifluorescence microscopy images of magnetically-activated SNAs and *in-situ* mNG expression in synthetic cells with and without exposure to an AMF. Synthetic cells (visualized through the encapsulated Texas-red dextran) that were not exposed to an AMF expressed minimal mNG, and their GFP channel fluorescence intensity was comparable to synthetic cells that contained PURExpress and the inactive DNA template only. Synthetic cells exposed to an AMF expressed mNG, as demonstrated by the increase in the green channel fluorescence, and were consistent with synthetic cells containing the full mNG template (dsDNA T7 promoter region present). Images are representative of n = 3 independent experiments. Scale bar = 20 μm. **(B)** Quantification of mNG expression in the individual GUVs using a circle detection-based image analysis script. Median fluorescence intensity (inactive template) = 0.00 grey units; median fluorescence intensity (SNA, no AMF) = 3.03 grey units; median fluorescence intensity (full template) = 17.47 grey units; and median fluorescence intensity (SNA, AMF) = 23.22 grey units. The box plot, notch and asterisk represent the interquartile range, mean and median fluorescence intensity, respectively. **(C)** Epifluorescence microscopy images of magnetically-activated SNAs and *in-situ* mNG expression in synthetic cells within a tissue phantom (black tube, see image). Synthetic cells that were not exposed to an AMF expressed minimal mNG. Whereas synthetic cells exposed to an AMF expressed mNG, unperturbed by the tissue phantom.

The ability for magnetic fields to deeply-penetrate bodily tissue confers significant advantages over previously published remote stimuli, predominantly light (*3, 36*). Moreover, magnetic hyperthermia is EMA/FDA approved for the treatment of glioblastomas and the resulting clinical formulation NanoTherm® (aminosilane-coated ferrofluid) is comparable to our central scaffold (*19*). To demonstrate superiority over the state-of-the-art and the applicability of magnetic fields as a deeply-penetrating external stimulus, we activated synthetic cells within a tissue-mimicking phantom (black tube) (Fig 4C). Tissue phantoms are synthetic materials that mimic the properties of biological tissues. In our case, we used a black tube as an optical tissue phantom that is impenetrable to light, the current state-of-the-art remote stimulus. As was previously observed, there was minimal leakage of mNG expression in the absence of an AMF (median fluorescence intensity = 5.06 grey units) (Fig 4C and Fig. S7). Astonishingly, the “on” state and the ability to control the magnetically-activated SNAs and *in-situ* mNG expression with exposure to an AMF was not perturbed by the tissue phantom (median fluorescence intensity = 24.55 grey units). The conservation of a high mNG fluorescence after exposure to an AMF in a tissue-mimicking phantom lays the foundations for synthetic cells as controllable drug delivery devices.

## Conclusions

We have described a robust strategy for controlling CFPS inside synthetic cells using magnetic hyperthermia, a clinically-approved and deeply tissue-penetrating stimulus, by generating magnetically-activated spherical nucleic acids. Beyond this, further nanoparticle engineering will allow for the exploration of the inherent properties of the core IONPs as contrast agents for magnetic resonance imaging (MRI), introducing the potential application of synthetic cells in theragnostics (*37*), and in magnetophoresis, controlling the movement of synthetic cells *in-vivo* (*38*). To our knowledge, this is the first example of controlling the activity of nucleic acids and/or CFPS with an AMF. The promising “off” state, showing negligible expression without the magnetic field, was only achieved with our newly developed SNA purification technique, which will prove essential to realising synthetic cells as targeted therapeutic modalities. Importantly, we have shown the selective control of CFPS in synthetic cells using magnetic field parameters (field strength, frequency and timeframe) compatible with the only clinically available magnetic hyperthermia system by MagForce, optimised to minimise eddy currents within the body and reduce patient discomfort (*19*). The ability to control the activity of synthetic cells within a tissue-mimicking phantom renders the developed technology advantageous over small-molecule or light activation, which are the current state-of-the-art. This selective regulation of biosynthesis *in-situ* demonstrates the remote control of synthetic cells using deeply tissue-penetrating magnetic fields and lays the foundations for the application of synthetic cells as smart delivery devices.

## Acknowledgments

We would like to thank Dr Gemma-Louise Davies and Prof Nguyen T. K. Thanh for their invaluable guidance and support on the nanoparticle synthesis and the application of magnetic hyperthermia. We would like to thank Prof Kylie Vincent and Prof Yujia Qing for their supervision support and input throughout the project. We would like to thank Dr Jack Howley for directing us to the thermal decomposition set-up and further Quentin Gueroult and the Goodwin lab for their help in acquiring the XRD data. E.P. is grateful to the EPSRC Centre for Doctoral Training in Inorganic Chemistry for Future Manufacturing (EP/S023828/1) for a studentship, generously supported by SCG Chemicals, Johnson Matthey, William Blythe, Siemens, Boron Specialties, Drochaid Research Services (DRS), Econic, High Force, OXECO, Oxford Instruments, and ISIS and Diamond facilities. M.J.B., A.A-S, G.M., and C.N. are supported by a Royal Society University Research Fellowship (URF\R\231007), the Biotechnology and Biological Sciences Research Council (BB/W011468/1 and BB/T008709/1), and the Engineering and Physical Sciences Research Council (EP/V030434/2 and EP/Y032675/1).

## Author contributions

E.P. and M.J.B. designed the project. E.P. designed, performed, and analysed the experiments, with contributions from all authors. E.P. and M.J.B. wrote the paper.

## Data availability

All the data generated in this study are available within the article, the supplementary information, and figures. Source data will be made available on Zenodo upon acceptance of the manuscript.

## Supplementary Materials

Materials and Methods, DNA sequences, Calculations, and Supplementary Figs. 1–7.

