## Supplementary Information for "Magnetic Activation of Spherical Nucleic Acids for the Remote Control of Synthetic Cells"

### Materials and Methods

#### Materials

All solvents and reagents were purchased from Sigma/Merck unless stated otherwise. Iron acetylacetonate was purchased from Fluorochem Ltd. Dibenzocyclooctyne-*N*-hydroxysuccinimidyl (DBCO-NHS) ester was purchased from Cambridge Bioscience Ltd. DreamTaq DNA polymerase master mix, TR-dextran (10 kDa) and 25 µL gene frames were purchased from Thermo Fisher Scientific. PURExpress, Lambda Exonuclease, Purple Loading Dye (6X) and RNA Gel Loading Dye (2X) was purchased from New England Biolabs. Egg PC was purchased from Avanti Polar Lipids. T7 promoter sequences (azide and non-modified) were synthesised by Integrated DNA Technologies.

#### Characterisation

Dynamic light scattering (DLS) was performed with a Zetasizer-Nano-ZS (Malvern Instruments, Ltd) at 25°C with nanoparticle solution at a concentration of 50 mg $L^{-1}$ to obtain the hydrodynamic diameter, polydispersity index and zeta potential.

X-ray diffraction (XRD) patterns were obtained using a Xcalibur PX Ultra diffractometer equipped with a CoKα radiation source (λ = 1.79 Å) operated at 40 mA. The crystallite diameter was evaluated by considering the X-ray wavelength, λ = 0.154 nm, with the crystallite-shape factor *K* set to 0.9, and the line broadening at the full half-width maximum of the most intense peak (in radians) and the Bragg angle (θ = 0.31 rad) (*1*).

$$D_{XRD} = \frac{K\lambda}{\beta\cos(\theta)}$$

Transmission electron microscope (TEM) images were captured using JEOL CX100 II TEM (JEOL, Ltd) at 100kV. Nanoparticle dispersions (0.1 mg $mL^{-1}$ in ethanol) were drop-casted onto carbon-coated copper grids and air-dried at room temperature. Particle size analysis was performed on >100 particles using ImageJ software.

The magnetic properties were characterised using a MPMS-5S superconducting quantum interference device (SQUID) magnetometer (Quantum Design, Ltd).

Alternating magnetic fields were generated using NAN201003 Magnetherm (Nanotherics Ltd).

#### Methods

**Synthesis of oleylamine-capped iron oxide nanoparticles**

Iron acetylacetonate (1.0595 g, 3 mmol) was dissolved in 15 mL of benzyl ether and 15 mL oleylamine. The reaction was evacuated and filled with argon five times (the vacuum was held for 5 min per cycle) and the reaction vessel was purged further with argon for 5 min to remove any remaining moisture. The solution was dehydrated at 110°C from room temperature under an argon atmosphere, before being heated to 293°C and aged at this temperature. The solution was magnetically stirred throughout the synthesis. Ethanol (40 mL) was subsequently added to the reaction mixture and the black material was precipitated and separated via centrifugation. The precipitate was dissolved in cyclohexane. Undispersed residue was removed by centrifugation. The product was precipitated with ethanol and centrifuged to remove the solvent, and re-dispersed into cyclohexane. The resulting oleylamine-capped iron oxide nanoparticles were stored at 4°C in cyclohexane (*2*, *3*).

**Silica encapsulation and amine modification of iron oxide nanoparticles**

IGEPAL-*co*-520 (0.5 mL, 0.5 g, 1 mmol) was dispersed in cyclohexane (11 mL, 101 mmol) and injected with oleylamine-capped iron oxide nanoparticles dispersed in cyclohexane (1 mL, 15 mg $mL^{-1}$). The dispersion was sonicated for 10 min at ambient conditions. Ammonium hydroxide 1M (0.13 mL, 0.13 mmol) was added to a magnetically stirred solution. The controlled addition of tetraethylorthosilicate (TEOS) (317 µL, 1.42 mmol) was performed using fractionated drop-method; where a known volume of TEOS (39.6 µL) was added every 3 hours. A total of three additions were carried out per day and reacted overnight, whereby the fourth addition was carried out 24 hours after the first addition and the seventh addition was carried out 48 hours after the first addition. Three hours after the eighth addition, (3-aminopropyl)triethoxysilane (APTES) (39.4 µL, 0.17 mmol) was added and the reaction was left overnight. The reaction was then diluted with ethyl acetate. Centrifugation was applied to pellet the nanoparticles and the solvent was removed. The nanoparticles were then dispersed in ethanol by sonication and diluted with ethyl acetate (3 parts) before pelleting by centrifugation and removing the solvent. This step was repeated twice more to remove the IGEPAL-*co*-520. The resulting silica-encapsulated iron oxide nanoparticles were then dispersed in ethanol (1 mL) to give a concentration of 15 mg/mL and stored at 4°C (*4*).

**Modification of amine-modified silica-encapsulated iron oxide nanoparticles**

Amine-modified silica-encapsulated iron oxide nanoparticles (300 µL, 5 mg/mL) were re-dispersed in 3-morpholinopropane-1-sulfonic acid (MOPS) buffer (0.1 M, pH 7.4). To this, DBCO-NHS ester (150 µL, 100 mM in dimethylformamide (DMF)) was added. The resulting dispersion was briefly sonicated and incubated overnight at room temperature in a ThermoMixer C with shaking at 1000 rpm. The DBCO-modified silica-encapsulated iron oxide nanoparticles were purified by centrifugation (13,300 rpm, 10 min), washing three times in ethanol, and stored in ethanol (5 mg/mL) at 4°C.

**Synthesis of spherical nucleic acids**

The 5′-azide-modified reverse compliment of the T7 promoter was annealed to the T7 promoter sequence in equimolar concentrations prior to heating at 95°C for 5 min (and cooling to room temperature over the course of 1 h) and analysis by polyacrylamide gel electrophoresis (PAGE). A native 16% PAGE was set-up at a 5 mL scale containing 40% acrylamide/bisacrylamide solution (2 mL), 10X TBE buffer (0.5 mL), water (2.5 mL), ammonium persulfate (40 µL, 10 w/v%) and TEMED (2.5 µL). The reaction was briefly vortexed and left to set for 30 min. The samples were prepared by mixing the DNA (180 ng, 10 µL) in Native Loading Dye (2X, 10 µL). Gels were run with 1X TBE buffer at 250 V for 60 min and then stained in 3X Gel Red for 10 min. Gels were imaged using Azure 200 Gel Imager (Azure Biosystems).

DBCO-modified silica-encapsulate iron oxide nanoparticles were dispersed in water. Sodium chloride was added (300 mM). To this, azide-modified dsT7 promoter was added. The reaction was then left overnight (*5*). The resulting T7 promoter-modified silica-encapsulated iron oxide nanoparticles (magnetically-activatable SNAs) were purified by centrifugation and agarose gel.

**Synthesis of the inactive mNG template**

DreamTaq MM (2X) (60 µL) was mixed with phosphorylated T7 forward primer or -T7 forward primer (2.4 µL, 25 µM), CT-rev or phosphorylated CT-rev respectively (2.4 µL, 25 µM), mNG PURExpress control template plasmid (*6*) (1.2 µL, 2 ng/µL) and water (54 µL). Reactions were cycled according to the following program using a ProFlex PCR System (ThermoFisher): 95°C for 60 s, 35 cycles (95°C for 30 s, 49°C for 30 s and 72°C for 65 s), 72°C for 10 min, 4°C HOLD. PCR products were validated by 1.5% TBE agarose gel electrophoresis and purified using a Monarch PCR & DNA Cleanup Kit NEB.

Double stranded mNG templates with and/or without the dsT7 promoter region (1000 ng, 10.66 µL) were diluted with water (11.34 µL). Lambda exonuclease (0.5 µL) and lambda exonuclease reaction buffer (10X, 2.5 µL) was added to a final volume of 25 µL. The reaction was allowed to proceed at 37°C for 30 min (template without the dsT7 promoter) and 35 min (template with the T7 promoter) before the lambda exonuclease was denatured at 70°C for 10 min. The crude reaction mixtures were combined and annealed at 95°C for 5 min (before slow cooling to room temperature). The resulting inactive mNG template was purified by Monarch PCR & DNA Cleanup Kit NEB and the purity was validated by 1.5% TBE agarose gel electrophoresis.

**Bulk cell-free protein synthesis of mNG protein**

Cell-free protein synthesis reactions (3 µL) were prepared with PURExpress Solution A (1.2 µL), PURExpress Solution B (0.9 µL) and supplemented with murine RNase inhibitor (0.075 U $\mu L^{-1}$). For the negative control, inactive mNG template (5 ng $\mu L^{-1}$, 0.30 µL) and water (0.52 µL) were added; for the positive control the full mNG template (with the dsT7 promoter present) (5 ng $\mu L^{-1}$, 0.30 µL) and water (0.52 µL) were added; and for the magnetically-activated SNA samples, inactive mNG template (5 ng $\mu L^{-1}$, 0.30 µL), magnetically-activatable SNAs (200 nM [as measured by calibration curve, see Fig. S4], 0.30 µL) and water (0.22 µL) were added. If necessary, the reactions were exposed to an alternating magnetic field (103.4 kHz, 30 mT) for 25 min and left for a further 25 min at room temperature. All reactions were then incubated in a thermal cycler at 25°C for 3 h and worked up with 17 µL of water. Reactions were transferred to Perkin Elmer 384-well black flat-bottom OptiPlates. Fluorescence was measured using a Tecan Spark fluorescence plate reader (Tecan Group, Ltd) (bandwidth, 5 nm; z position, 17,764 µm; ExmNG, 495 nm; EmmNG, 517 nm).

**Assembly of mNG and magnetic-activation of synthetic cells**

Egg PC dissolved in chloroform (25 mg $mL^{-1}$, 150 µL) was transferred to a 2 mL glass vial (per two conditions). The vials were tilted at 45° and rotated slowly while held under $N_2$ flow to distribute the lipids evenly up the walls. Vials containing lipid films were held under a vacuum for 3 h to remove residual chloroform. Mineral oil (filtered through 0.22 µm PES membrane) was added to Egg PC films (0.674 g) to give a final concentration of 5 mg $mL^{-1}$ Egg PC. Vials were vortexed for 1 min and then incubated in the oven at 80 °C for 10 min with the lids removed. Lids were reapplied to vials and sealed tightly using Teflon tape and parafilm, then vortexed aggressively for 1 min and sonicated in a sonication bath for 1 h at 50°C. Lipid-containing oil was stored at room temperature overnight and vortexed, then sonicated for 10 min at room temperature immediately before use. A total of 250 µL of 5 mg $mL^{-1}$ Egg PC in mineral oil was transferred to 1.5 mL centrifuge tubes and placed on ice. A total of 10 µL inner solution (PURExpress containing 5 ng $\mu L^{-1}$ inactive mNG template or full mNG template, 400 nM magnetically-activatable SNAs (as measured by calibration curve, see Fig. S4), 1.0 U μL−1 murine RNase inhibitor, 25 μM TR-dextran (10 kDa) and 200 mM sucrose) was added into the chilled lipid-containing oil, ensuring the tip was constantly moved through the lipid-containing oil to disperse the inner solution. Tubes were passed across a centrifuge rack once using light pressure to form cloudy water-in-oil (W/O) emulsions. Meanwhile, 100 μL of lipid-containing oil was layered on top of 250 μL of chilled outer solution (50 mM HEPES, 400 mM potassium glutamate and 200 mM glucose (pH 7.6)) and placed at room temperature. W/O emulsions were then added on top of this oil phase, and this tube was left at room temperature for 1 min. Centrifuge tubes containing the W/O emulsion above the outer solution were centrifuged at 16,000 x g, 4 °C, for 30 min. After centrifugation, the oil phase and ~200 μL of the outer solution were removed from the tube and discarded. Using a fresh tip, ~10 μL of the remaining outer solution was ejected against the GUV pellet to displace it from the tube, and the intact GUV pellet was transferred to a new tube containing 250 μL outer solution. The pellet was subsequently resuspended by gently pipetting up and down. Vesicles were centrifuged at 10,000 x g, 4 °C for 10 min, then the outer solution was removed and the pellets were gently resuspended in 25 μL of fresh outer solution (*6*).

**mNG expression in synthetic cells**

To assess mNG expression from inside the synthetic cells, the GUV pellets were resuspended in 25 μL outer solution and placed in colourless or black (tissue phantom) microcentrifuge tubes. If necessary, the synthetic cells were exposed to an alternating magnetic field (103.4 kHz, 30 mT) for 10 min and left for a further 10 min at room temperature. All reactions were then incubated in a thermal cycler at 25°C for 2.5 h and worked up with 25 μL of outer solution (50 mM HEPES, 400 mM potassium glutamate and 200 mM glucose (pH 7.6)). Fluorescence microscopy was performed using an EVOS® FL Fluorescence Microscope using a ×100 oil immersion objective lens. GUVs were imaged using the brightfield, TXR and GFP filters.

**Image processing**

TXR and GFP channel brightness was normalised across all images within a single experiment, and then the separate channels were saved as individual PNG files. All images corresponding to a single sample were stored within the same directory. "Background" images were created by manually selecting vesicle-free regions of microscopy images (one from each sample within the experiment) and measuring the mean pixel intensity. PNG files were input into the vesicle_analysis script as described in previous literature (see https://zenodo.org/record/7729425 for script) (*6*).

### DNA Sequences

| DNA name | DNA sequence | Modification |
|---|---|---|
| T7 Promoter Sequence (Sense/Top Strand) | GAAATTAATACGACTCACTATAG | |
| T7 Promoter Sequence (Antisense/ Bottom Strand) | CTATAGTGAGTCGTATTAATTTC | 5′-azide C6 modifier |
| -T7 Forward Primer | GTTTAACTTTAAGAAGGAGGTATACATATGGTGAG | |
| Phosphorylated CT-Rev | GATATAGTTCCTCCTTTCAG | 5′-phosphorylated |
| Phosphorylated T7 Forward Primer | GAAATTAATACGACTCACTATAGGGTCTAG | 5′-phosphorylated |
| CT-Rev | GATATAGTTCCTCCTTTCAG | |
| mNG Linear Template | gaaattaatacgactcactatagggtctagaaataattttgtttaactttaagaaggaggtatac<br>atATGGTGAGCAAAGGCGAAGAGGATAATATGGCAAGC<br>CTGCCTGCAACACATGAACTGCATATTTTTGGTAGCAT<br>TAACGGCGTGGATTTTGATATGGTTGGTCAAGGCACCG<br>GTAATCCGAATGATGGTTATGAAGAACTGAATCTGAA<br>AAGCACCAAAGGCGATCTGCAGTTTAGCCCGTGGATTC<br>TGGTTCCGCATATTGGTTATGGTTTTCATCAGTATCTGC<br>CGTATCCGGATGGTATGAGCCCGTTTCAGGCAGCAATG<br>GTTGATGGTAGCGGTTATCAGGTTCATCGTACCATGCA<br>GTTTGAAGATGGTGCAAGCCTGACCGTTAATTATCGTT<br>ATACCTATGAAGGCAGCCACATTAAAGGTGAAGCACA<br>GGTTAAAGGTACAGGTTTTCCGGCAGATGGTCCGGTTA<br>TGACCAATAGTCTGACCGCAGCAGATTGGTGTCGTAGC<br>AAAAAAACCTATCCGAACGATAAAACCATCATCAGCA<br>CCTTCAAATGGTCATATACCACCGGCAATGGTAAACGT<br>TATCGTAGCACCGCACGTACCACCTATACCTTTGCAAA<br>ACCGATGGCAGCAAACTATCTGAAAAATCAGCCGATG<br>TATGTGTTTCGCAAAACGGAACTGAAACATTCCAAAAC<br>CGAGCTGAACTTTAAAGAATGGCAGAAAGCATTTACC<br>GATGTGATGGGTATGGATGAGCTGTACAAATAATGAgg<br>atcccgggaattctcgagtaaggttaacctgcaggaggcctttaattaaggtggtgcggcc<br>gcgctagcggtcccgggggatcgatccggctgctaacaaagcccgaaaggaagctgagt<br>tggctgctgccaccgctgagcaataactagcataaccccttggggcctctaaacgggtctt<br>gaggggttttttgctgaaaggaggaactatatc | |

### Calculations

#### Determination of nanoparticle concentration

The nanoparticle concentration (particles $mL^{-1}$) was calculated according previous literature(*7*) by approximating each individual IONP@$SiO_2$ as spherical in shape. The mass of the core-shell IONP@$SiO_2$ ($m_{c\text{-}s}$) was estimated using the radii of both the core IONPs ($r_c$) and core-shell IONPs@$SiO_2$ ($r_{core\text{-}shell}$) from TEM analysis, and the known densities of silica ($\rho = 2.20$ g $cm^{-3}$) and magnetite ($\rho = 5.24$ g $cm^{-3}$) (Eq. S1–3). The concentration of the core-shell IONP@$SiO_2$ ($N_{core\text{-}shell}$) was calculated from the mass concentration of nanoparticles in mg $mL^{-1}$ ($M_C$) and the mass of the individual core-shell IONP@$SiO_2$ ($m_{core\text{-}shell}$) (Eq. S4).

$$m_{core} = \frac{3}{4}\pi\, r_{core}^{3}\, \rho_{magnetite} \qquad \text{Eq. S1}$$

$$m_{shell} = \frac{3}{4}\pi\left(r_{core-shell}^{3} - r_{core}^{3}\right)\rho_{silica} \qquad \text{Eq. S2}$$

$$m_{core-shell} = m_{core} + m_{shell} \qquad \text{Eq. S3}$$

$$N_{core-shell} = \frac{M_C}{m_{core-shell}} \qquad \text{Eq. S4}$$

#### Determination of DBCO loading

The DBCO concentration (c) was calculated using the Beer Lambert Law (Eq. S5) from the absorbance (A) of the DBCO at 309 nm (taken on a NanoPhotometer C40 UV/Vis Spectrophotometer (Implen)), the molar extinction coefficient (ε) of DBCO at $\varepsilon = 12{,}000\ M^{-1}cm^{-1}$ (309 nm) (*8*), and a path length (l) of 0.01 cm.

$$A = \varepsilon c l \qquad \text{Eq. S5}$$

The DBCO concentration (N) was then converted to moles and Avogadro's number ($N_A$) was used to convert to number of DBCO molecules in a known volume (Eq. S6). The number of known DBCO molecules was divided by the number of nanoparticles present (calculated as described previously) to gain the number of DBCO molecules per nanoparticle.

$$N = N_A \times mol \qquad \text{Eq. S6}$$

#### Determination of DNA loading

DNA concentration on the SNAs was determined by denaturing the attached dsDNA at 95°C for 5 min in RNA Gel Loading Dye (2X) and comparing the concentration (and further moles) of DNA released to a calibration curve of known DNA concentrations (Fig S4). Following this, the concentration of DNA strands was calculated using Avogadro's number ($N_A$) (Eq. S6) and divided by the number of nanoparticles present.

### Supplementary Figures

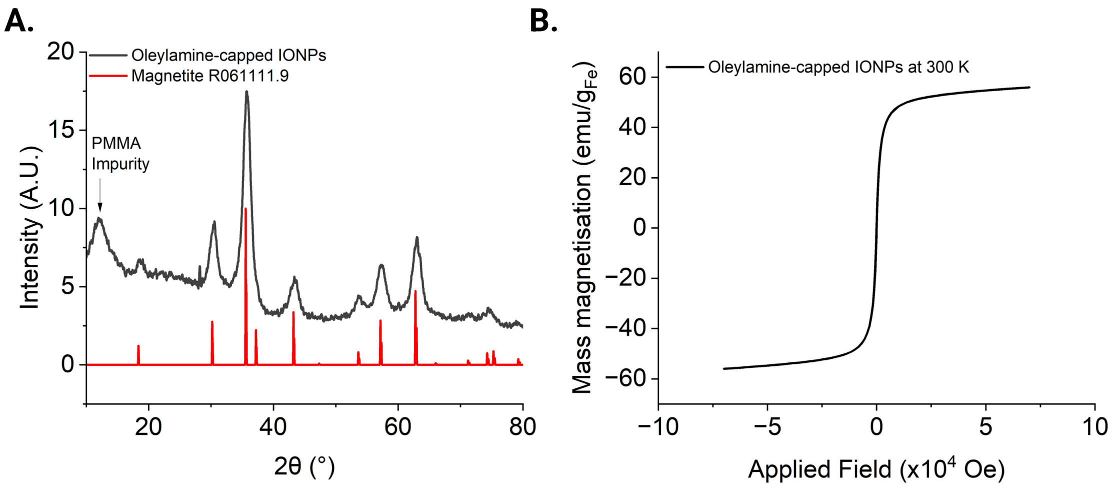

**Fig. S1. Characterisation of oleylamine-capped IONPs. (A)** XRD pattern of oleylamine-capped IONPs indexed against magnetite (RUFF ID:R061111). **(B)** Hysteresis curve of oleylamine-capped IONPs measured at 300 K.

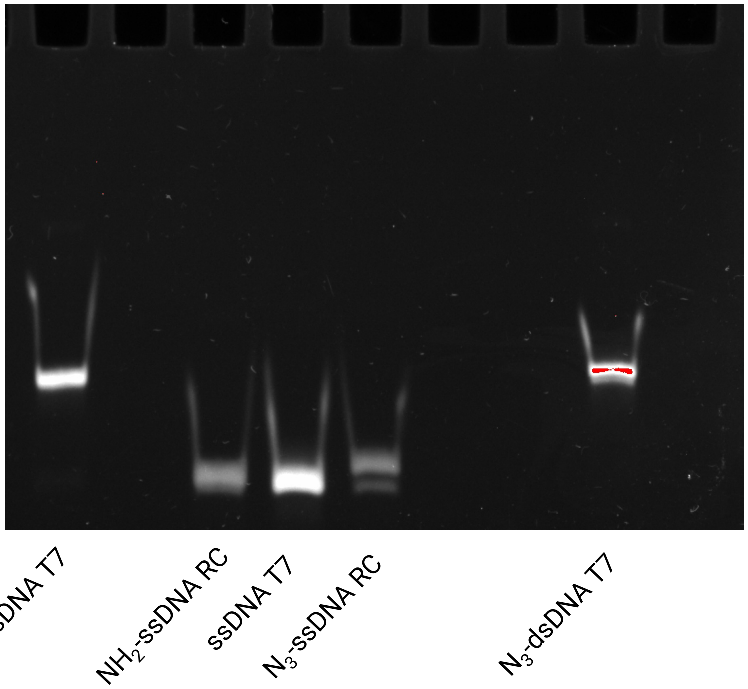

**Fig. S2. PAGE of DNA annealing.** Native 16% poly(acrylamide) gel electrophoresis tracking the successful annealing of the T7 promoter antisense strand with ($N_3$-ssDNA RC) and without ($NH_2$-ssDNA RC) the $N_3$ click handle to the T7 promoter sense strand (ssDNA T7), and the resulting dsDNA with ($N_3$-dsDNA T7) and without ($NH_2$-dsDNA T7) the $N_3$ click handle. The dsDNA is of higher molecular mass and so travels slower down the gel in the direction of the electric current (top to bottom of the gel).

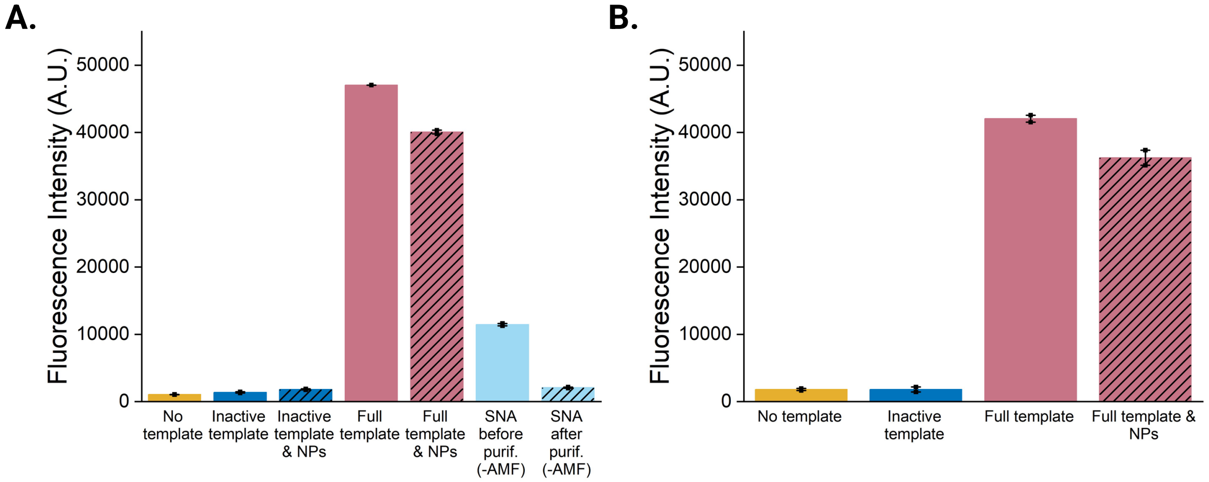

**Fig. S3. Effect of SNA purification by agarose gel and presence of nanoparticles in cell-free expression. (A)** In vitro transcription of the Broccoli RNA aptamer that after expression binds to the small molecule DFHBI and fluoresces. RNA expression was recovered with the full Broccoli template compared to the inactive Broccoli template (without the ds T7 promoter) and the RNA expression and fluorescence intensity was not perturbed by the addition of $NH_2$-modified IONPs@$SiO_2$. The tight "off" state of the SNA (-AMF), comparable to the inactive template (negative control), was only achieved after agarose purification of the SNAs. **(B)** Cell-free protein synthesis of mNG in the presence of $NH_2$-modified IONPs@$SiO_2$, showing negligible inhibition of mNG expression and fluorescence intensity compared with the full mNG template (dsDNA T7 promoter region present) without the nanoparticles present.

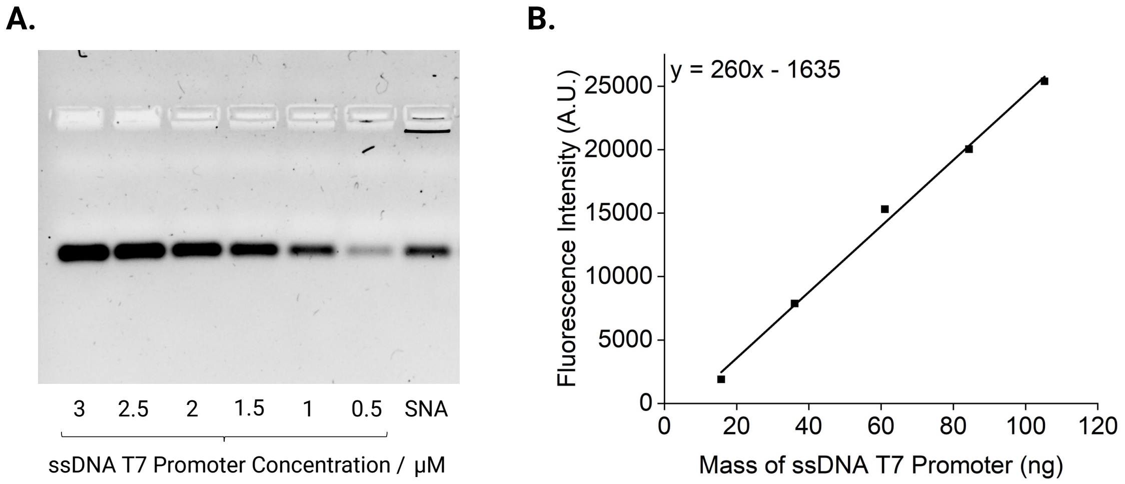

**Fig. S4. Calibration curve to determine SNA T7 promoter loading. (A)** Agarose gel of known concentrations of the T7 promoter sequence and a known volume of SNA heated at 95°C in urea-containing dye to release the bound T7 promoter. **(B)** Calibration curve of the Gel Red fluorescence intensities (determined by ImageJ) against the known T7 promoter concentration, the linear regression (fitted in Origin) was used to extrapolate the concentration of bound T7 promoter on the SNA.

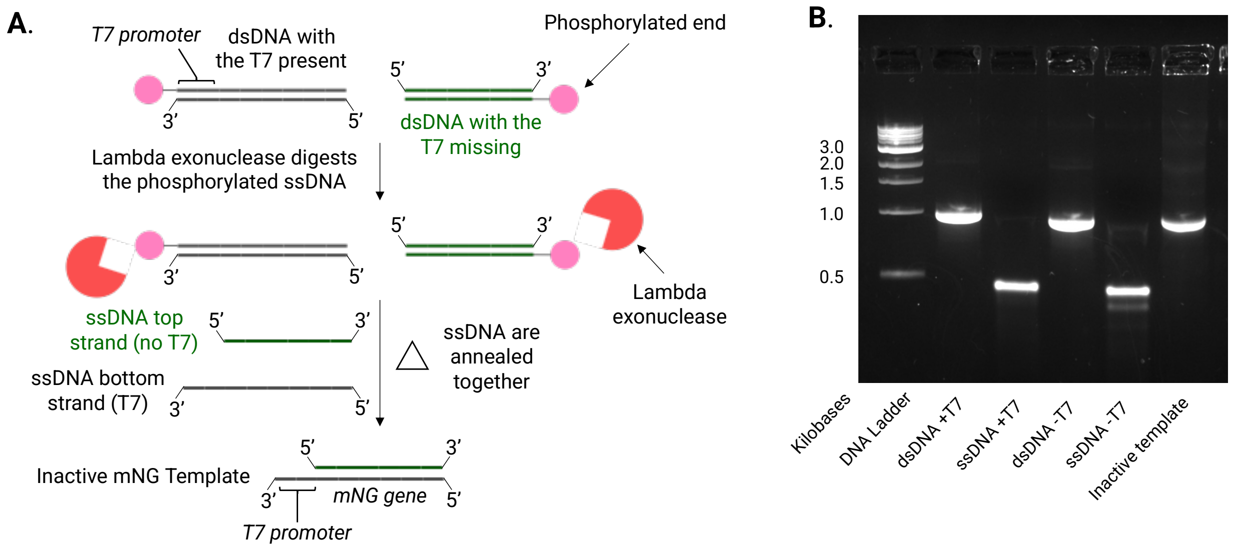

**Fig. S5. Synthesis of the inactive mNG template. (A)** Schematic showing the synthesis of dsDNA with and without the T7 promoter present and with the sense and antisense strand, respectively, phosphorylated, and the subsequent digestion of the phosphorylated strands by lambda exonuclease prior to annealing to create the inactive mNG template. Note: promoter (23 base pairs) and gene (~700 base pairs) length are not to scale. **(B)** Native 1.5% agarose gel quantifying the synthesis of the inactive template with the dsDNA with (dsDNA +T7) and without (dsDNA -T7) the T7 promoter region present, its digestion to the ssDNA antisense strand with the T7 promoter present (ssDNA +T7) and the ssDNA sense strand without the T7 promoter (ssDNA -T7), prior to the annealing and formation of the inactive template (inactive template).

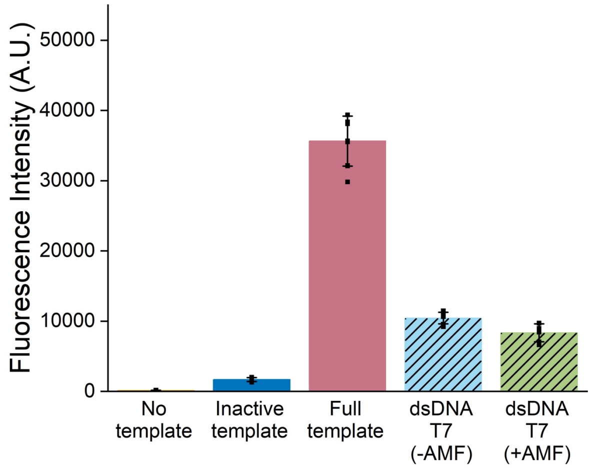

**Fig. S6. Cell-free protein synthesis with an alternating magnetic field and without magnetic nanoparticles.** Cell-free protein synthesis of mNG in the presence of dsT7 promoter with and without an AMF. There is no increase in mNG expression and fluorescence intensity with exposure of the dsT7 promoter to an AMF, showing that the denaturing of the dsDNA, release of the T7 promoter and recovery of the inactive mNG template is not due to the heat generated from the solenoid coil.

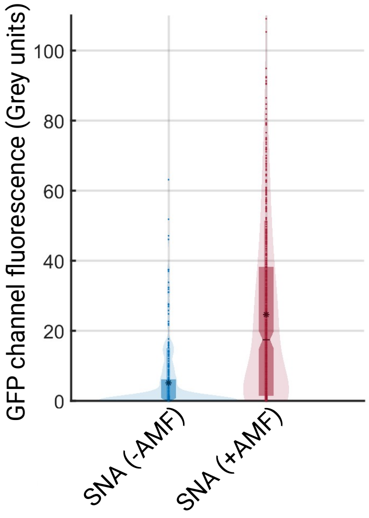

**Fig. S7. Quantification of mNG expression in a tissue phantom.** mNG expression in the individual GUVs using a circle detection-based image analysis script. Median fluorescence intensity (SNA, no AMF) = 5.06 grey units and median fluorescence intensity (SNA, AMF) = 24.55 grey units. The box plot, notch and asterisk represent the interquartile range, mean and median fluorescence intensity respectively.
